# Comparing vibrotactile stimulation to combined visual and auditory stimulation for 40 Hz gamma entrainment

**DOI:** 10.1101/2025.03.14.639935

**Authors:** Yeongdae Kim, Ayush Choudhary, Hyeonseok Kim, Peter Teale, Brice McConnell, Mazen Al Borno

**Affiliations:** Computer Science and Engineering, University of Colorado at Denver | Anschutz Medical Campus, Aurora, Colorado, United States; Division of Child and Adolescent Psychiatry, Cincinnati Children’s Hospital Medical Center, Cincinnati, Ohio, United States; Department of Neurology, University of Colorado at Denver | Anschutz Medical Campus, Aurora, Colorado, United States

**Keywords:** non-invasive stimulation, vibrotactile stimulation, Alzheimer’s disease

## Abstract

There is significant interest in combined visual and auditory stimulation to entrain 40 Hz gamma oscillations for the treatment of Alzheimer’s disease and other neurological conditions such as stroke and insomnia. In this work, we compared another sensory modality—vibrotactile stimulation delivered with a glove—to visual and auditory stimulation in 15 healthy participants in terms of electroencephalogram (EEG) responses and subjective experience. We found that vibrotactile stimulation from the glove could evoke 40 Hz EEG responses in the central, frontal and, to a lesser extent, occipital cortices. We also observed distinct patterns of functional connectivity between the two stimulation modalities. Participants preferred the vibrotactile stimulation over the visual and auditory stimulation. Our study supports future investigations on vibrotactile stimulation for the treatment of neurological conditions.

## Introduction

Alzheimer’s disease (AD) is a neurodegenerative condition without an effective pharmacotherapeutic treatment that is estimated to affect 5.7 million in the USA [1] and 30-35 million worldwide [2]. Brain oscillations are disrupted in AD and other neurological conditions such as Parkinson’s disease and epilepsy. There is ongoing research in developing non-invasive methods to restore physiological brain oscillations for these diseases [3]. Recent studies found that combined visual and auditory stimulation (VAS) at the gamma frequency of 40 Hz reduced pathologies associated with AD and improved cognitive function in mouse models of neurodegeneration [4,5]. More specifically, stimulation at the 40 Hz gamma frequency, but not other frequencies, reduced the accumulation of Beta-amyloid plaques and phosphorylated tau protein that characterize AD [6]. Furthermore, six weeks of one hour of daily VAS reduced neuronal and synaptic loss, which are hallmarks of neurodegenerative diseases. VAS had more beneficial effects than either visual or auditory stimulation alone, including decreasing amyloid in medial prefrontal cortex [5]. Another modality, namely somatosensory stimulation based on whole-body vibrotactile stimulation, also decreased brain pathology associated with AD in the primary somatosensory cortex and primary motor cortex [7]. Other techniques such as vagal nerve stimulation can attenuate hippocampal amyloid load and improve cognition and memory in certain mouse models [8].

Based on these studies in mouse models, pilot studies in humans with AD are underway with VAS. Recent clinical studies show promising preliminary evidence that prolonged stimulation positively impacts neural activity and immune cells [9,10], as well as improves sleep quality and activities of daily living in patients with prodromal AD [11]. In a randomized controlled trial, AD patients in the treatment group received VAS from neurostimulation glasses for an hour per day for six months, which resulted in reduced white matter and myelin loss compared to the sham group, as assessed with magnetic resonance imaging [12]. In addition to AD, gamma oscillations induced by 40 Hz VAS can rescue functional synaptic plasticity after stroke in mice [13] and promote sleep in children with insomnia [14].

Clinical application of VTS remains limited, but EEG-based studies have shown that 20 Hz VTS induces stable responses in the somatosensory cortex over extended periods without signs of habituation [15], and other studies have also demonstrated reliable entrainment at 40 Hz in healthy participants [16,17]. This evidence indicates that VTS can provide a reliable means of driving brain oscillation activity. Though VAS showed clinical feasibility in AD patients, daily exposure to one hour of VAS presents a substantial challenge, even among patients actively seeking treatment as the flashes of light (i.e., light flickering) can limit their vision and prevent their activities of daily living, and the auditory tone can be difficult to tolerate. For example, Hajós et al. [18] reported that 18.5 % of participants declined participation during screening and 28% dropped out during treatment. Meanwhile, VTS has the practical advantage of being applicable during activities of daily living. Most human studies of VTS have concentrated on active applications such as brain-computer interfaces, rather than on reactive entrainment responses [17,19]. As a result, evidence regarding its tolerability and neurophysiological effects in humans remains preliminary. These considerations highlight the need for studies directly comparing VAS and VTS in humans as a stimulation approach prior to clinical application.

In this study, we investigated VTS as a potential sensory modality for 40 Hz putative gamma entrainment. The mechanical vibration was applied on the back of the hand, which produces a sensation like a phone vibrating. We evaluated EEG responses from two minutes of VAS and VTS to 15 healthy participants in terms of response locations and changes in brain connectivity. In addition, participants verbally answered a questionnaire to assess the tolerability of the different stimulation modalities.

## Materials & Methods

Each participant first underwent VAS, rested for 5 minutes, and then underwent VTS. Each stimulation lasted for 2 minutes. We recorded EEG responses during the stimulation, as well as 2 minutes before (pre-stimulation) and after the stimulation (post-stimulation), resulting in a total of 6 minutes of recordings per sensory modality. Participants were seated 2 feet away from the device and asked to keep their eyes open during VAS. For VTS, participants wore right-hand gloves and were asked to either keep their eyes open or closed for the whole experiment. Lastly, we conducted additional EEG recordings with VAS for 5 subjects on a different day. In this session, we used an eight-channel EEG headset (the OpenBCI Ultracortex ‘Mark IV’ EEG Headset) for a 250 Hz sampling rate and computed event-related potentials from VAS.

### Participants

15 healthy adults (4 females, age M = 28.6, SD = 6.7, Range 19-39) were recruited from the University of Colorado Denver community. A priori power analysis was conducted using G*Power for a within-subject design, with α = .05 and a power of .80, which indicated that a sample size of 15 participants was sufficient to detect a large effect size (Cohen’s d ≥ .8) on 40 Hz power, consistent with established methodological reporting guidelines in psychophysiological research [20] and effect sizes reported in prior work with 40 Hz visual stimulation [21]. Consequently, while this study was adequately powered to identify robust, large-scale EEG differences, findings regarding smaller effect sizes should be interpreted as exploratory.

With both verbal and written explanations of the study requirements, all participants provided written informed consent, as approved by the Institutional Review Board at the University of Colorado Denver (COMIRB 23-0783). All methods were performed in accordance with relevant guidelines and regulations.

### Data Collection Methods

EEG signals were recorded using 16 dry electrodes from the OpenBCI Ultracortex ‘Mark IV’ EEG Headset, structured in a 10-20 configuration (see Fig. 2A). We used two reference nodes on the earlobes. The measured impedances ranged between 15 kΩ and 80 kΩ for all channels, which are below acceptable thresholds [22,23]. The data sampling rate was 125 Hz. For one subject, whose head size did not fit the dry electrode headset, gel-type neuroelectric sensors were used instead.

**Figure 1.**
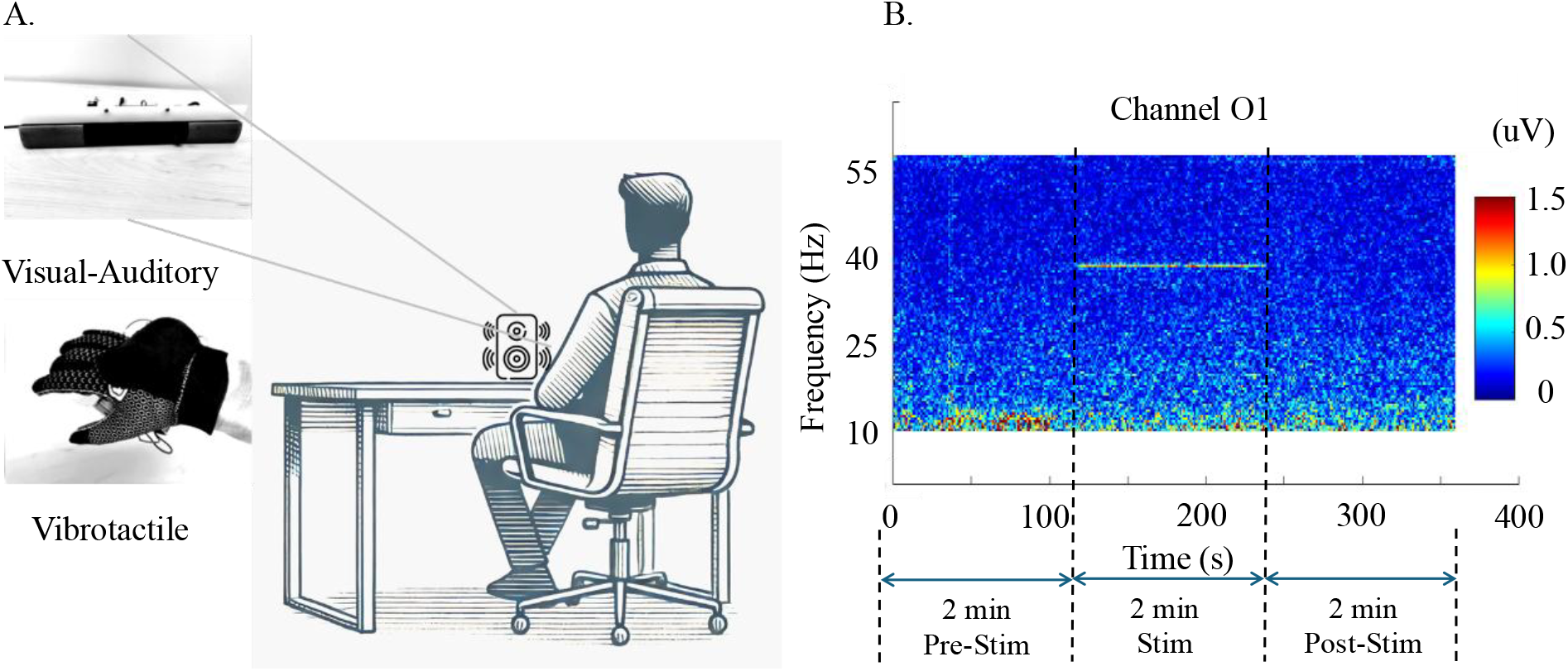
**A**. Illustration of the experimental session. Participants, sitting on a chair, receive VAS from a device on the desk. After a 5-minutes rest period, participants receive VTS from a glove worn on their right hand. **B**. Time-frequency spectrogram of channel ‘O1’ from VAS. The stimulation is applied for two minutes, between 120-240 s. The stimulation causes a peak in power (uV) at the 40 Hz frequency.

**Figure 2.**
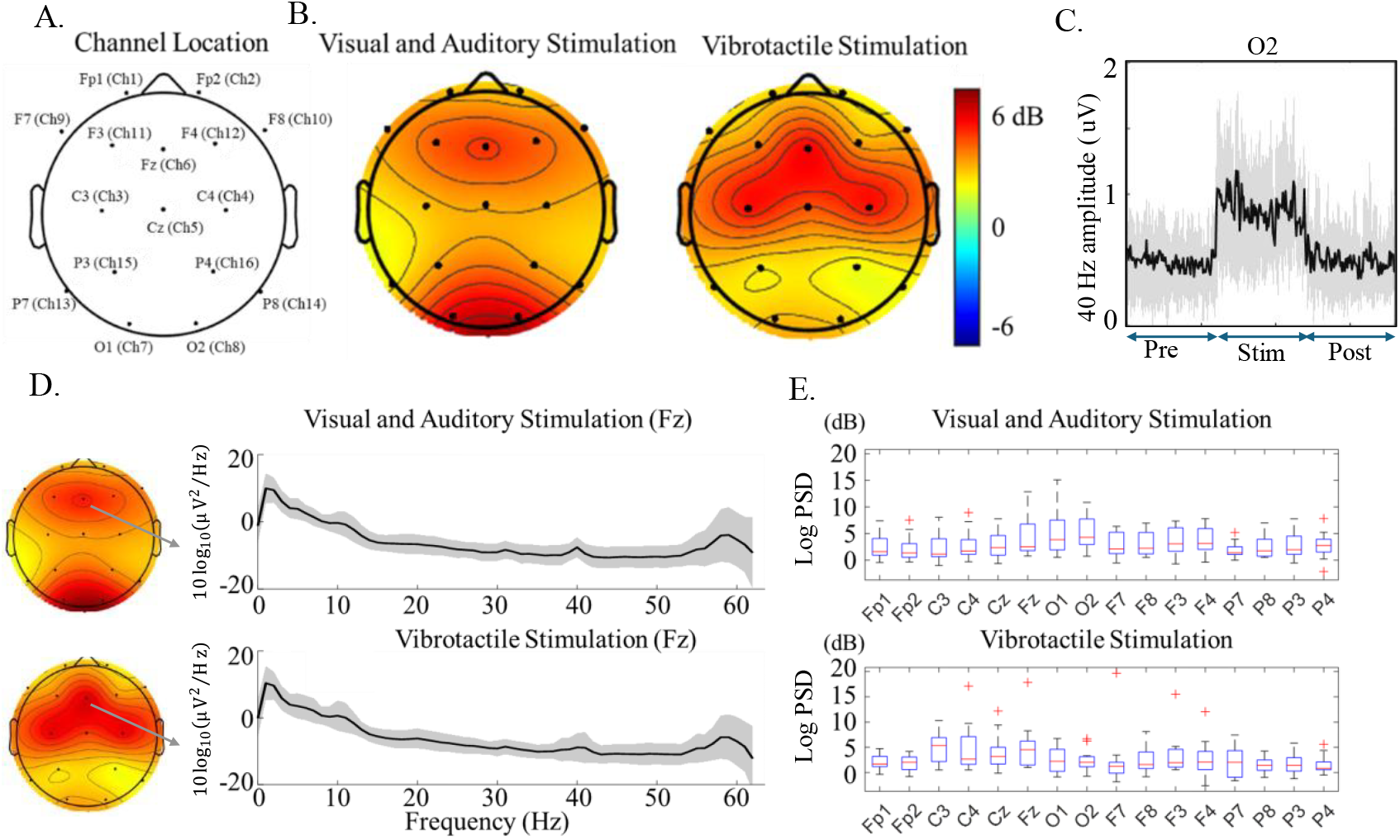
**A**. Sensor locations on the scalp in our EEG study with 16 channels. **B**. Respective putative brainwave entrainment for each stimulation modality. The size of the putative entrainment is represented by the relative increase in intensity compared to the pre-stimulation period, displayed on a dB scale. The colors represent the relative intensity of stimulation onset putative brain entrainment from the 40 Hz stimulation frequency compared to the pre-stimulation period. **C**. Time series of the 40 Hz response during the experiment for sensor O2 from VAS. The black solid line represents the average across all participants, and the gray shading indicates on standard deviation from the mean. **D**. Log power spectral density of the pre-dominantly entrained channel Fz for the two stimulation modalities. **E**. Boxplots of channel-specific brainwave putative entrainment (dB) for VAS (top) and VTS (bottom).

#### VTS apparatus

Because of the concentration of sensory endings and receptors on the hand [24], we applied a glove-based VTS. We used an eccentric rotating motor (ERM; Yooptop 2300 RPM 12V DC Motor) to produce the desired 40 Hz frequency using a constant voltage (3.65 V). We used a linear voltage regulator (LM337T), a dual differential comparator, a TO-220 heat sink, and multiple resistors to ensure a constant voltage supply or shut down the motor when the input was insufficient. The whole circuit was soldered onto a PCB board. The glove was comprised of the PCB board, an ERM motor, and a 9V lithium-ion battery. The PCB board and the motor are inside a 3-D printed case, sewn onto the glove to stimulate the backside of the hand. We chose the backside of the hand because it causes less interference with activities of daily living than the palm. Furthermore, our pilot testing with 5 participants indicated that it was at least as effective as the palm. When constructing the glove apparatus, a stable frequency of 40 Hz was confirmed in two subjects in pilot testing. However, during the actual experiments, we observed a frequency variation of approximately 5%, ranging from 38 to 42 Hz. This deviation was caused by vibrations in the subject’s hand and the motor itself attached to the glove. In future work, we plan to use an accelerometer to monitor the frequency in real-time and adjust the voltage accordingly to address this limitation in accuracy with VTS.

#### VAS apparatus

For VAS, a sound bar ‘AS500’ from Dell with a top-mounted light source was used. The light source consisted of 36 Rextin LEDs of 5050 LED (5,0 × 5.0 mm, with three diodes per package), arranged in a rectangular bar with dimensions of 50 cm in length and 1.2 cm in width. The audio input signal was split between the sound bar and a circuit to synchronously drive the LEDs. The LED driver input was high-pass filtered at 340 Hz and applied to an inverting op-amp with a gain of 20, which overloaded the amplifier and generated a negative output pulse to trigger a 555 timer. The timer was set to produce a 12.5 ms (50% duty cycle) positive pulse, which was applied to a transistor switch in series with the LED power source. The audio stimulator pulse was a 1ms-long 10 kHz tone [5], delivered by a computer audio file and presented at 58 dB in amplitude.

### EEG signal preprocessing

The EEG signals were imported into MATLAB 2022a (The MathWorks, Inc.) using EEGLAB v2024.0 [25]. Sensor locations were assigned based on default boundary element model from DIPFIT v5.4 [26,27], and these standardized locations were later utilized for signal source analysis. EEG signals were filtered using a 60 Hz line noise filter [28] and finite impulse response band-pass filter between 1-60 Hz. Re-referencing was performed to the average reference, with a zero-padded channel included to ensure the data retained full rank [29]. Using the re-referenced EEG data, we computed 1-second quantized spectrograms and excluded data that showed atypical uniform increase across all frequencies, which can be caused by sensor dislocation, muscle activity or eye blinking [30].

Following noise removal, independent component analysis (ICA) was deployed using AMICA [31], an adaptive mixture ICA function to decompose the EEG data into mutually independent components (ICs) with corresponding topographic projections on the scalp sensors. To systematically isolate and remove artifacts, the decomposed ICs were automatically classified using the ICLabel plugin [32]. ICs with a ‘Muscle’ or ‘Eye’ probability greater than 50% were excluded, resulting in an average of 3.33±1.54 rejected ICs per participant. The signal source dipole locations of the remaining ICs were then computed using DIPFIT, which localized each IC within the Montreal neurological institute template [33]. Each dipole location, initially a single 2×2×2 mm voxel, was transformed into a dipole density sphere using 3-dimensional Gaussian smoothing with a 20-mm full width at half maximum. This transformation mitigated inter-subject variability in dipole locations and determined distributed activation patterns across specific anatomical regions. The resulting dipole density was mapped onto anatomical regions using an automated anatomical labeling system, enabling consistent localization across subjects and group-level analysis [34].

### EEG signal analysis

We analyzed the relative intensity changes between the stimulation and pre-stimulation periods to examine whether and where brainwave entrainment occurred. Narrow-band signal-to noise ratio was computed as the ratio of power at the stimulation frequency to the median power of its neighboring frequency bins within ±0.75 Hz, 0.25 Hz steps, excluding the target bin, following frequency-tagging practice [35,36]. We used the median instead of the mean to obtain a robust estimate against outlier bins. Individual entrainment was defined by a robust z score at the target stimulation relative to the neighboring-bin distribution, with responders identified at z ≥ 1.96 and a power increase of at least 3 dB compared to the pre-stimulation period. Condition effects on response intensity were quantified as relative dB between the stimulation and pre-stimulation periods and tested at the group level with paired t-tests.

We also investigated the EEG connectivity shifts due to stimulation. In channel-based functional connectivity analysis, coherence between EEG channel pairs was quantified across canonical frequency bands. All coherence values were Fisher z-transformed, and statistical significance was evaluated using sign-flip permutation tests with multiple-comparison correction by the Benjamini-Hochberg false discovery rate [37]. To detect effects that were not apparent at the channel-level, we computed effective connectivity across ICs using renormalized partial directed coherence (RPDC) with time-frequency decomposition. This analysis was conducted at a sampling rate of 125 Hz, and 45 frequency bins logarithmically spaced between 2 and 50 Hz with single-window analysis for the pre-, during and post-stimulation periods. This process produced a connectivity matrix of varying number of IC × IC for each subject per three periods. Lastly, weak family-wise error rate (FWER) control was applied to analyze the statistical significance of changes in time–frequency decomposed RPDC across the pre-, during and post-stimulation periods.

## Results

We compared the effects of the different stimulation modalities on the neural circuit based on the putative entrainment location on the scalp near the 40 Hz stimulation frequency. For VAS, 14 out of 15 participants exhibited significant putative entrainment in at least one of the 16 channels. Similarly, for VTS, 13 out of 15 participants showed putative entrainment in at least one channel. In Fig. 2, we illustrate the relative intensity of stimulation onset brain entrainment at the 40 Hz frequency compared to the pre-stimulation period, averaged over all subjects for VAS and VTS. As shown in Fig. 2B and 2E, the primary response of the VAS was the occipital area with a 5.3 ± 3.0 dB intensity increase at the O2 sensor. In Fig. 2C, we show the 40 Hz response in O2 during the pre-stimulation, stimulation and post-stimulation periods. The other main response occurred in the frontal cortices adjacent to channel Fz with a 4.2 ± 3.4 dB intensity increase. VTS showed the highest 40 Hz stimulation-locked response on the frontal-central region with an average 4.8 ± 4.3 dB intensity on sensor Fz. The second most putatively entrained sensor is C3 in the left hemisphere, as participants wore gloves on their right hand. Nearly half of the participants (n=7) demonstrated strong 40 Hz stimulation-locked responses in the occipital and adjacent parietal regions with VTS (with more than 3 dB intensity increase). We excluded the possibility that this putative entrainment was due to the mechanical vibration of the motor (between 38 to 42 Hz), as we verified that the accelerometer (placed near the occipital regions) did not exhibit a peak near 40 Hz frequency. In Fig. 2D, we show the power spectral density mean and standard deviation on a predominantly entrained channel (Fz). The response amplitude to the VTS was larger than that of the VAS. We did not vary the stimulation intensity in this study. Variations in the motor frequency (between 38-42 Hz due to the glove apparatus) resulted in a more widespread response near the 40 Hz target with VTS, averaging at 39.97 ± 1.32 Hz over the 15 participants. The 40-Hz putative entrainment was not observed in the post-stimulation condition for VAS and VTS.

In channel-based coherence, the only significant difference observed across the three periods (pre-, during, and post-stimulation) was between the stimulation and pre-stimulation periods in the gamma band with VAS (Fig. 3A). Specifically, the parietal-frontal pair (F4-P4) channel-based raw coherence decreased from 0.187 ± 0.115 to 0.165 ± 0.110 (ΔC = −0.022), with a permutation-based *p* < .0001 that survived BH-FDR correction (q < .05). Furthermore, using weak FWER control, we identified statistically significant connectivity shifts for both VAS and VTS. During VAS, connectivity from the parietal regions (specifically the postcentral gyrus and the precuneus) to the frontal regions decreased in the beta band (13-30 Hz) compared to the pre-stimulation period (Fig. 4A-B). A similar pattern of decreased connectivity was observed after stimulation, though in a different anatomical location, specifically from the left calcarine to left cingulum (Fig. 4C). This effect was frequency-specific and consistently observed across multiple anterior-posterior IC pairs. Following VTS, we observed a significant increase in connectivity between the left superior medial frontal gyrus and the left orbital frontal region compared to the pre-stimulation period (see Fig. 4D). This change in connectivity was observed in the gamma band, which corresponds to the 40 Hz stimulation frequency.

**Figure 3.**
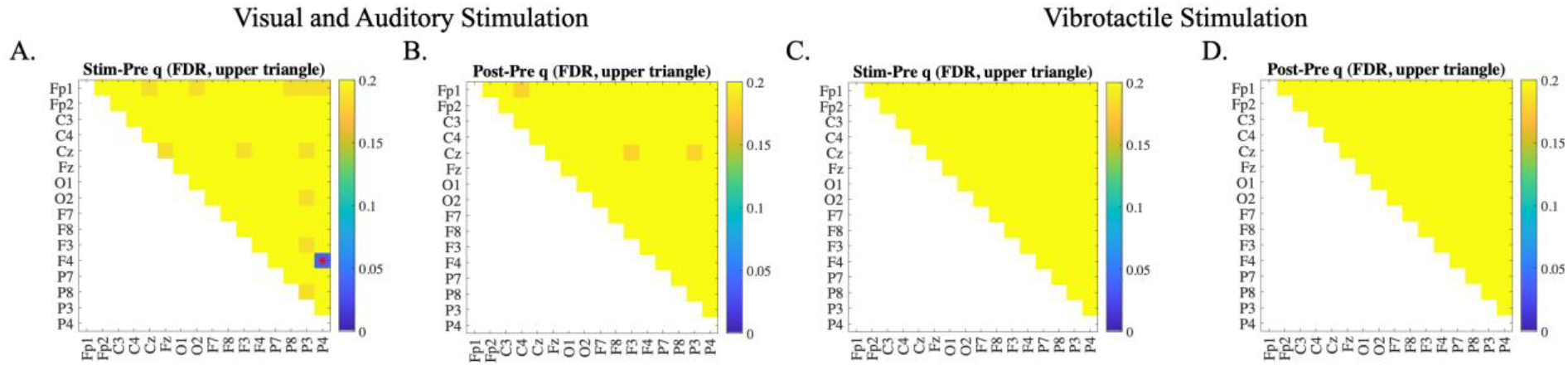
Channel-based coherence comparison between the stimulation and pre-stimulation periods in the gamma band, evaluated with Benjamini-Hochberg FDR correction (q values). The red dot highlights the only channel pair that reached statistical significance (F4-P4 in A). No statistically significant differences were observed in B, C and D. **A**. Coherence comparison between the stimulation and pre-stimulation period in the VAS condition. **B**. Coherence comparison between the post-stimulation and pre-stimulation period in the VAS condition. **C**. Coherence comparison between the stimulation and pre-stimulation period in the VTS condition. **D**. Coherence comparison between the post-stimulation and pre-stimulation period in the VTS condition.

**Figure 4.**
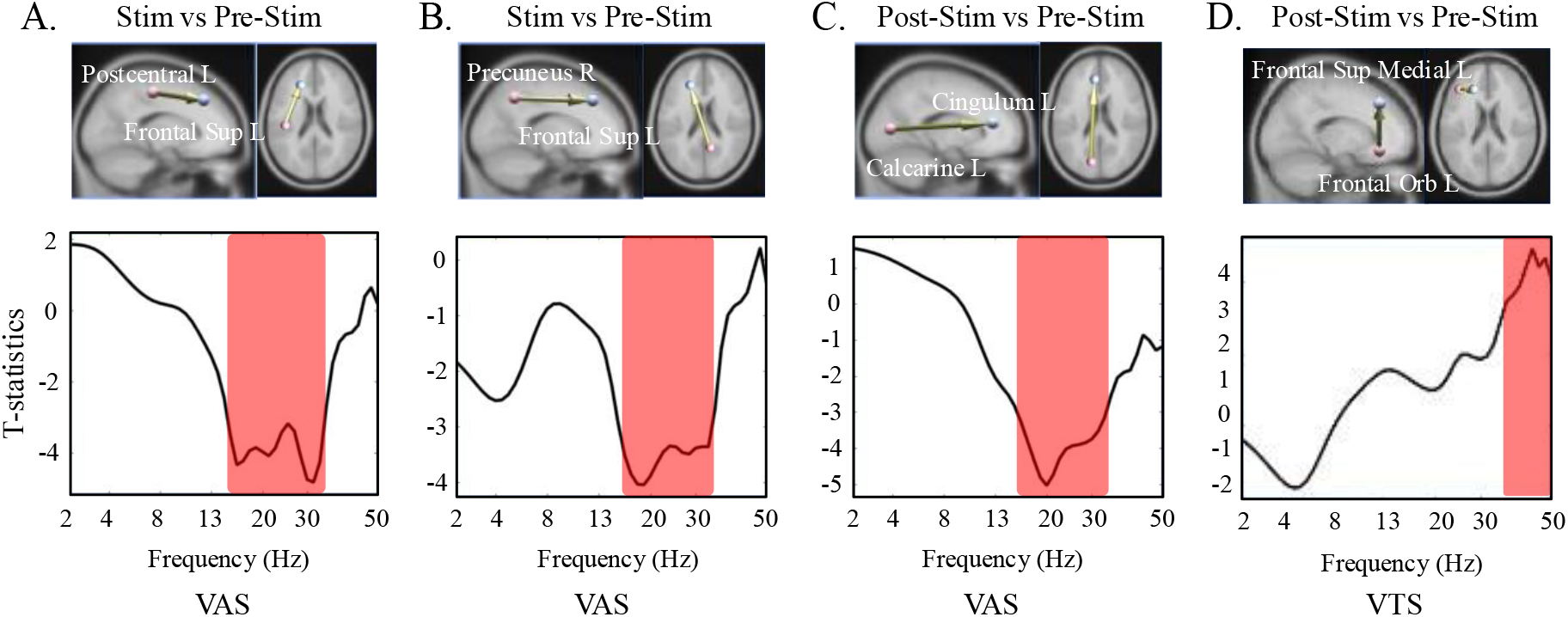
**A**. Connectivity shift measures with VAS (comparing the stimulation and pre-stimulation periods) in the 13-30 Hz frequency band between the left postcentral gyrus and left superior frontal cortex, depicted in a connectome connectivity map. The red-highlighted regions of the frequencies indicate t-statistics with an absolute value greater than 3 (|t(14)| > 3, p < .01). **B**. Same as A except the regions are the right precuneus and the left superior frontal cortex. **C**. Same as A except the comparisons are between the post-stimulation and pre-stimulation periods, and the regions are the left calcarine and left cingulum. **D**. Same as C except that it is for VTS, and the regions are the left superior medial frontal gyrus and the left orbital frontal region.

In Supplementary Fig. 1, we computed the event-related potentials for one subject during 2 minutes of VAS (26 samples per 100 ms epoch). We observed that among the seven EEG channels fluctuating at 40 Hz, two channels from the occipital region showed more than double the amplitude and were antiphase to the other channels. We found that 2 out of 5 participants had this approximate π phase difference between the occipital and the other entrained cortices. We could not reliably estimate the phase difference in the remaining subjects because the sample rate was still too low.

We conducted verbal interviews with participants after undergoing the experiment. Their assessments of each stimulation modality are summarized in Table 1. Participants were asked to evaluate how long they could receive the stimulation daily for months and rate their tolerance on a scale from 0 to 10, where 0 indicated extreme difficulty in tolerating the stimulation and 10 indicated no difficulty. The tolerance scores for VAS and VTS were 6.2 ± 2.4 and 8.8 ± 1.3, respectively, showing higher tolerance for the VTS (*p* = .00027). All participants indicated higher daily duration tolerance for VTS (67.7 min ± 75.8) compared to VAS (11.5 min ± 15.4; *p* = .0049). Participants reported experiencing eyesight blurring from the visual stimulation, finding the auditory stimulation to be the most irritating sensory modality, and feeling bored during VAS. The consensus was that VTS is a more tolerable stimulation modality than VAS, causing less discomfort.

**Table 1.** Participants verbal responses to the stimulation modality.

| Stimulation Type | Visual and Auditory | Vibrotactile | P-value |
| --- | --- | --- | --- |
| Tolerance | 6.2 ± 2.4 | 8.8 ± 1.3 | < .01 |
| (0: intolerable / 10: tolerable) |  |  |  |
| Daily duration tolerance (minutes) | 11.5 ± 15.4 | 67.7 ± 75.8 | < .01 |
| Preferred modality (15 subjects) | 1/15 | 14/15 |  |

## Discussion

Non-invasive 40 Hz stimulation has recently received considerable attention because of promising experiments conducted on mouse models of neurodegeneration [4,5] and stroke [13]. Several studies are now underway in AD patients to assess the feasibility and safety of VAS and to assess the effects of the stimulation when applied over long durations on the neural circuitry [9–11]. In this study, we examined EEG responses from healthy participants to VAS and VTS and gathered participants’ feedback on their preferred stimulation modality.

Our experiments confirmed near 40 Hz brainwave responses with VAS and VTS. Teplan et al. [38] demonstrated short-lived brainwave entrainment in the occipital and frontal cortical regions across multiple frequencies (2, 4, 9, and 17 Hz). We observed similar patterns in these regions with 40 Hz VAS. In addition to the O1-O2 channels, VAS induced F3, Fz, and F4—regions associated with the dorsolateral and medial frontal cortices—indicating the involvement of attention and sensory processing in response to the combined stimuli. In contrast, VTS primarily induced C3, Cz, and C4 in the sensorimotor cortex, associated with processing tactile and proprioceptive information. A prior study found that VTS decreased brain pathology in these regions in mouse models of neurodegeneration [7]. VTS had smaller responses in the frontal cortices than VAS, perhaps due to the reduced attentional involvement demands of the stimulation. Furthermore, VTS exhibited strong occipital responses in seven out of fifteen participants, consistent with prior work showing the activation of occipital cortex during tactile processing [39,40]. Similarly, VAS also stimulated the somatosensory cortices (see Fig. 2A), which may encode visual information in working memory [41]. Our work shows that a cost-effective off-the-shelf ERM motor can reliably produce EEG responses near 40 Hz, whereas recent work was unsuccessful at this task [42]. This discrepancy may be due to differences in hardware considerations or device design. Nevertheless, given the absence of a dedicated sham or alternative-frequency control condition, we cannot rule out the possibility that these observed EEG dynamics partly reflect modality-specific evoked responses rather than pure cortical entrainment [43,44].

During VAS, we observed a distinct reduction in long-range connectivity. Specifically, pathway-relevant connectivity from the occipital area to the frontal area decreased in the beta band ([45]; see Fig. 4A-C). Furthermore, our channel-based coherence analysis revealed a significant reduction in gamma-band coupling between posterior and anterior sites (P4-F4) during stimulation. These findings suggest that VAS may induce a distinct, immediate modulatory effect, potentially by prioritizing sensory-driven activity that transiently suppresses long-range posterior-to-anterior communication.

In contrast, VTS did not induce significant connectivity change during the stimulation period, possibly indicating that vibrotactile stimulation requires a longer integration window or operates through different neural pathways than visual-auditory inputs. VTS elicited a significant increase in interconnectivity within the anterior frontal region in the gamma band during the post-stimulation period. This highlights a possible mechanistic difference: while may VAS induce rapid, transient network reconfiguration (reduced connectivity) between posterior and anterior regions, VTS appears to promote delayed or sustained local integration within the anterior frontal cortex. The absence of immediate locking in VTS, followed by a post-stimulation increase in the gamma band, suggests that VTS could potentially be associated with a cumulative electrophysiological change, though the definitive underlying neuroplastic mechanisms remain to be validated with a sham control and increased electrode density. Thus, VTS might be more advantageous for applications requiring sustained cortical excitability rather than immediate state changes. Because our analysis was performed with only 16 EEG electrodes, future work with high-density EEG could improve the spatial resolution of our findings. A limitation of our study is that we did not include a sham or different stimulation frequency. For this reason, we cannot assess whether the observed EEG changes are specific to the 40 Hz VAS or VTS or reflect more general effects of sensory stimulation or attention, especially on a connectivity basis. Future studies could incorporate appropriate control conditions to better isolate frequency-specific and modality-specific effects.

Prior work has shown that VAS is more effective than visual or auditory stimulation alone in reducing AD pathologies [5]. The brain adjusts for the differences in transmission and sensory processing times when integrating different sensory modalities [46]. Teplan et al. [47] investigated VAS and observed a stabilized angular difference after a certain period of sustained stimulation. We also found a phase difference between the auditory and visual 40 Hz stimulation-locked responses (see Supplementary Fig. 1). Future studies should investigate the optimal lags (if any) between sensory modalities to avoid negative interference at 40 Hz and improve treatment outcomes, which will be especially important as technologies with more sensory modalities are developed.

Based on the participants’ preliminary feedback on the stimulation, all but one participant preferred the VTS modality. The participant who preferred VAS also noted that the VTS could be easier to tolerate if applied daily. However, the fixed order of conditions (VAS preceding VTS in the study) may have introduced an order effect, potentially involving subjective factors reported during post-experimental interviews, such as a cumulative fatigue or a sense of relief once the initial stimulation was completed. Consequently, given this non-counterbalanced design and the short stimulation duration, our observations regarding participant preference and tolerability should be regarded as preliminary. Future studies should employ randomized or counterbalanced designs to control for potential order effects. We confirmed that the physiological baseline (pre-stimulation resting state) remained consistent between the VAS and VTS conditions, as there were no statistically significant differences (p > .05) in RPDC values or 40 Hz power. Since the stimulation would likely require daily sessions over months or even years, treatments can be burdensome for individuals. Therefore, VTS represents a potential alternative sensory modality to VAS, warranting further investigation into whether it could offer greater comfort and mobility, in addition to reducing limitations on social interactions and engagement with the external environment. As our study observations were based on short-term exposures (2-minute stimulation periods), the tolerability of extended daily use remains to be tested in future longitudinal studies. Moreover, for individuals with reflex epilepsy, such as photosensitive occipital lobe epilepsy, non-visual stimulus options may minimize risk. The incorporation of other stimulation modalities could help address the diverse needs of patients and enhance treatment outcomes. Extending treatment duration and improving compliance through such options may also be critical for promoting neural plasticity and supporting recovery.

## Conclusion

We confirmed that both VAS and VTS evoked near 40 Hz gamma oscillations in the sensory and frontal cortices, while producing distinct patterns of functional connectivity. Specifically, VAS was associated with an immediate and transient reduction in long-range coherence between posterior and anterior regions. In contrast, VTS preferentially enhanced local intra-frontal gamma connectivity during the post-stimulation period, suggesting a potential sustained after-effect and a different mode of network engagement.

Although our study was limited to a fixed stimulation intensity and short-term exposure to the stimulation, it provides a proof-of-concept for modality-specific gamma modulation. Our participant population were healthy young adults, which may limit generalizability to elderly or Alzheimer’s disease populations. Future studies could explore optimized stimulation parameters (such as frequencies and intensities), longer intervention periods, and the inclusion of clinical populations—such as those with Alzheimer’s disease—to establish the therapeutic potential of these differing stimulation paradigms.

## Supporting information

manuscript_images

## Acknowledgements

We are grateful to the Coleman Institute for Cognitive Disabilities and ARIAD CU Denver for funding this work.

## Data Availability

All the de-identified data from our study and code are available at https://doi.org/10.5281/zenodo.15008933

## Code Availability

Data analysis was performed using the open-source EEGLAB toolbox and its associated publicly available plugins. No bespoke custom code or novel algorithms were generated for this study.

## Supplementary Material

**Supplementary Figure 1.**
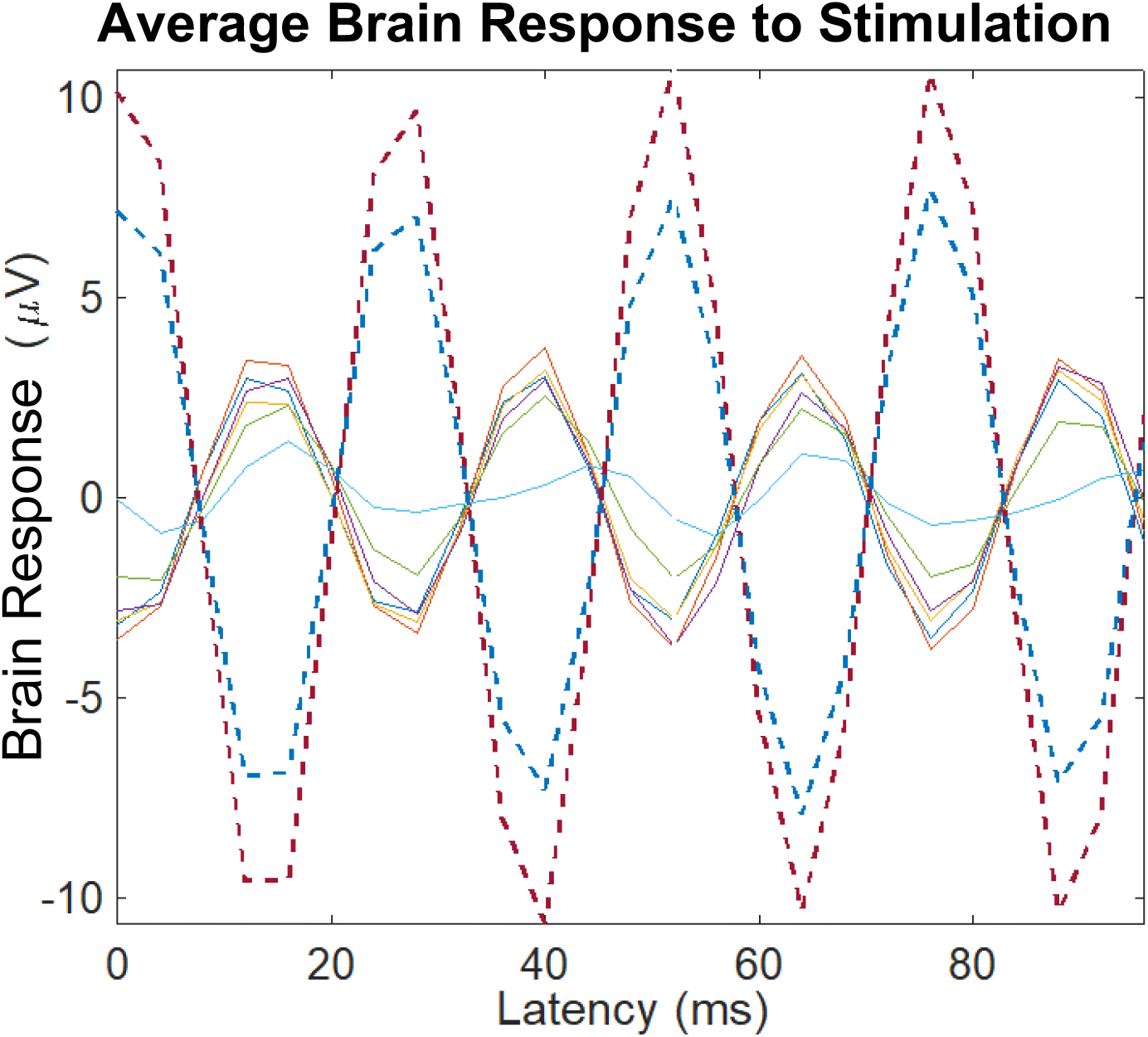
Event-related potentials for one subject during VAS. Solid lines represent the prefrontal, frontal and central cortex and dotted lines represent the occipital cortex.

