## Supplementary material for "Comparing vibrotactile stimulation to combined visual and auditory stimulation for 40 Hz gamma entrainment": manuscript_images: Figure 1.pdf

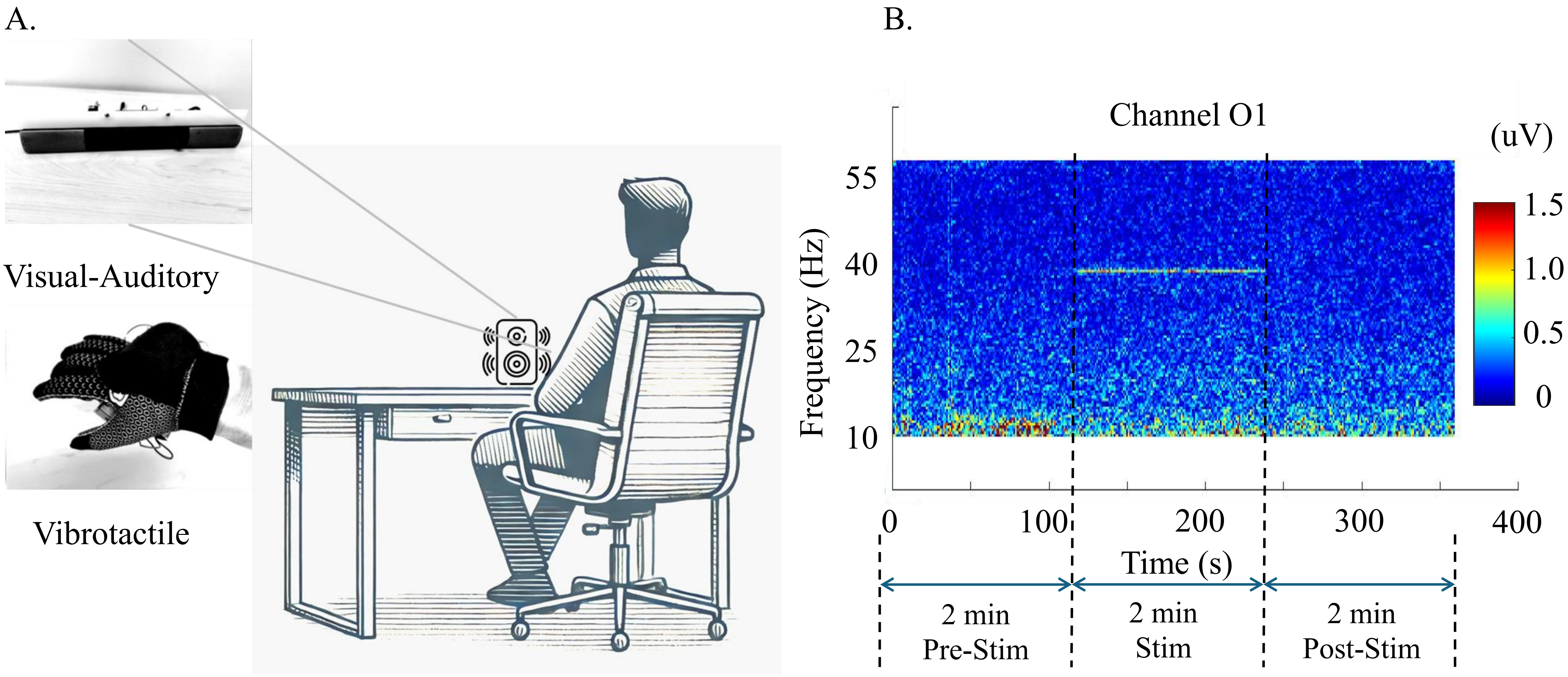


**Figure 1. A.** Illustration of the experimental session. Participants, sitting on a chair, receive VAS from a device on the desk. After a 5-minutes rest period, participants receive VTS from a glove worn on their right hand. **B.** Time-frequency spectrogram of channel 'O1' from VAS. The stimulation is applied for two minutes, between 120-240 s. The stimulation causes a peak in power (uV) at the 40 Hz frequency.
