## Supplementary material for "Comparing vibrotactile stimulation to combined visual and auditory stimulation for 40 Hz gamma entrainment": manuscript_images: Figure 2.pdf

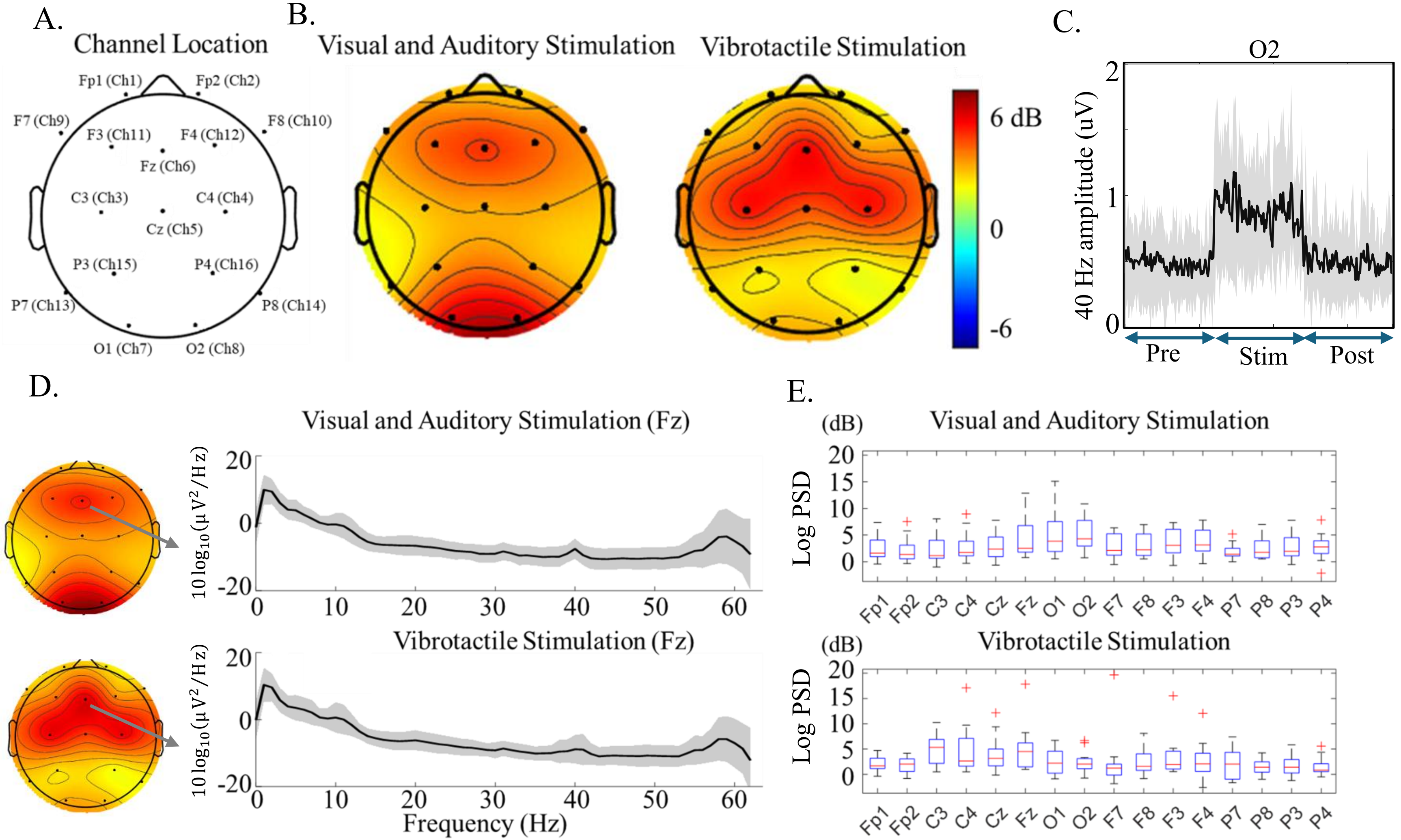


**Figure 2. A.** Sensor locations on the scalp in our EEG study with 16 channels. **B.** Respective putative brainwave entrainment for each stimulation modality. The size of the putative entrainment is represented by the relative increase in intensity compared to the pre-stimulation period, displayed on a dB scale. The colors represent the relative intensity of stimulation onset putative brain entrainment from the 40 Hz stimulation frequency compared to the pre-stimulation period. **C.** Time series of the 40 Hz response during the experiment for sensor O2 from VAS. The black solid line represents the average across all participants, and the gray shading indicates on standard deviation from the mean. **D.** Log power spectral density of the pre-dominantly entrained channel Fz for the two stimulation modalities. **E.** Boxplots of channel-specific brainwave putative entrainment (dB) for VAS (top) and VTS (bottom).
