## Supplementary material for "Comparing vibrotactile stimulation to combined visual and auditory stimulation for 40 Hz gamma entrainment": manuscript_images: Figure 3.pdf

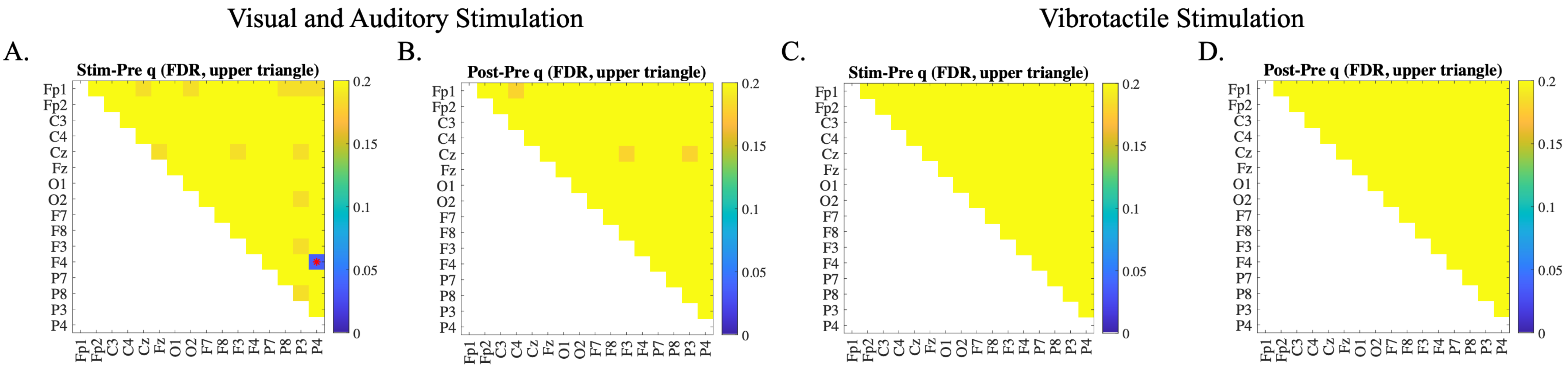


**Figure 3.** Channel-based coherence comparison between the stimulation and pre-stimulation periods in the gamma band, evaluated with Benjamini-Hochberg FDR correction (q values). The red dot highlights the only channel pair that reached statistical significance (F4-P4 in A). No statistically significant differences were observed in B, C and D. **A.** Coherence comparison between the stimulation and pre-stimulation period in the VAS condition. **B.** Coherence comparison between the post-stimulation and pre-stimulation period in the VAS condition. **C.** Coherence comparison between the stimulation and pre-stimulation period in the VTS condition. **D.** Coherence comparison between the post-stimulation and pre-stimulation period in the VTS condition.
