## Supplementary material for "Comparing vibrotactile stimulation to combined visual and auditory stimulation for 40 Hz gamma entrainment": manuscript_images: Figure 4.pdf

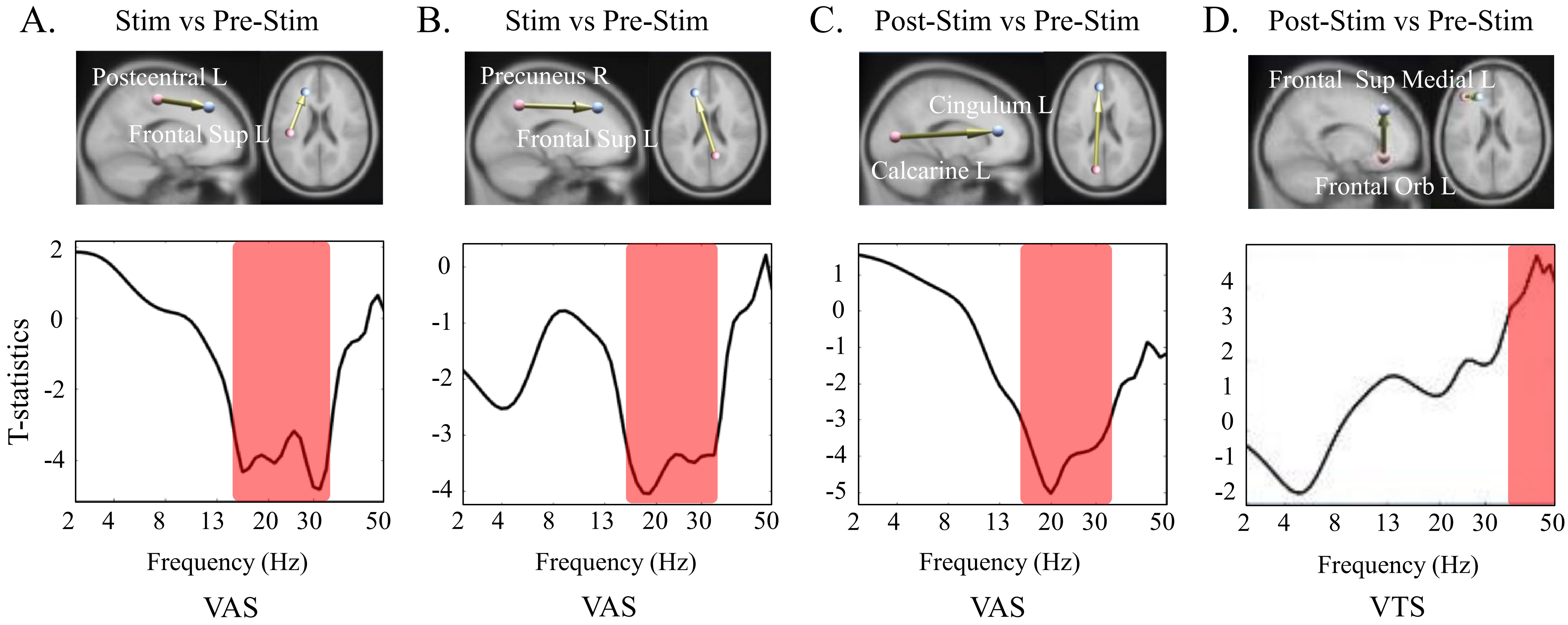


**Figure 4. A.** Connectivity shift measures with VAS (comparing the stimulation and pre-stimulation periods) in the 13-30 Hz frequency band between the left postcentral gyrus and left superior frontal cortex, depicted in a connectome connectivity map. The red-highlighted regions of the frequencies indicate t-statistics with an absolute value greater than 3 ($|t(14)| > 3$, $p < .01$). **B.** Same as A except the regions are the right precuneus and the left superior frontal cortex. **C.** Same as A except the comparisons are between the post-stimulation and pre-stimulation periods, and the regions are the left calcarine and left cingulum. **D.** Same as C except that it is for VTS, and the regions are the left superior medial frontal gyrus and the left orbital frontal region.
