## Supplementary material for "Comparing vibrotactile stimulation to combined visual and auditory stimulation for 40 Hz gamma entrainment": manuscript_images: Final_Manuscript.pdf

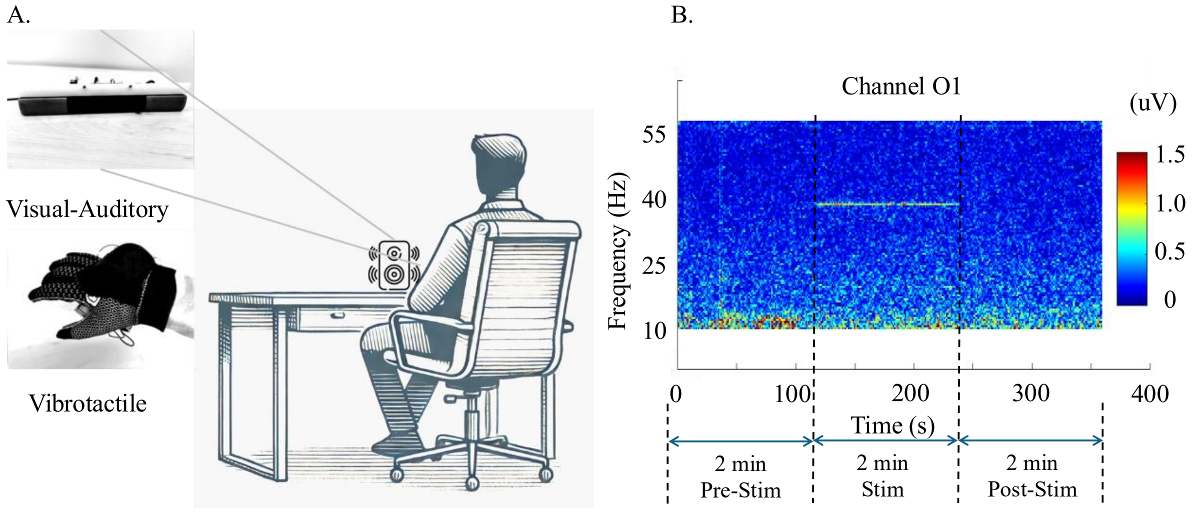


**Figure 1. A.** Illustration of the experimental session. Participants, sitting on a chair, receive VAS from a device on the desk. After a 5-minutes rest period, participants receive VTS from a glove worn on their right hand. **B.** Time-frequency spectrogram of channel 'O1' from VAS. The stimulation is applied for two minutes, between 120-240 s. The stimulation causes a peak in power (uV) at the 40 Hz frequency.

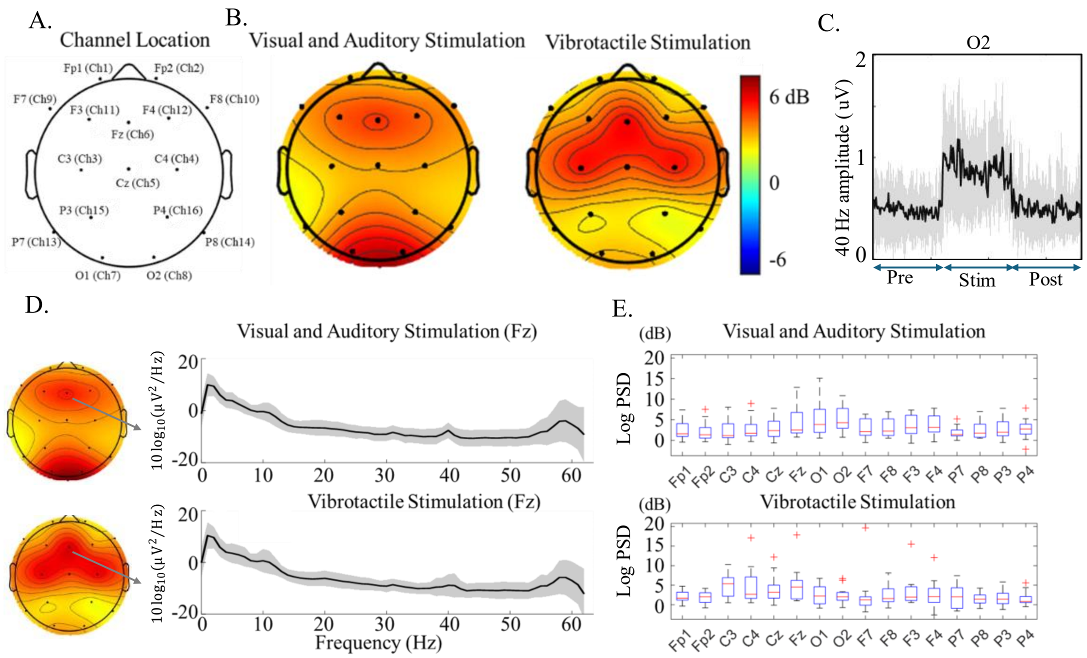


**Figure 2. A**. Sensor locations on the scalp in our EEG study with 16 channels. **B.** Respective putative brainwave entrainment for each stimulation modality. The size of the putative entrainment is represented by the relative increase in intensity compared to the pre-stimulation period, displayed on a dB scale. The colors represent the relative intensity of stimulation onset putative brain entrainment from the 40 Hz stimulation frequency compared to the pre-stimulation period. **C.** Time series of the 40 Hz response during the experiment for sensor O2 from VAS. The black solid line represents the average across all participants, and the gray shading indicates on standard deviation from the mean. **D.** Log power spectral density of the pre-dominantly entrained channel Fz for the two stimulation modalities. **E.** Boxplots of channel-specific brainwave putative entrainment (dB) for VAS (top) and VTS (bottom).

In channel-based coherence, the only significant difference observed across the three periods (pre-, during, and post-stimulation) was between the stimulation and pre-stimulation periods in the gamma band with VAS (Fig. 3A). Specifically, the parietal-frontal pair (F4-P4) channel-based raw coherence decreased from 0.187 ± 0.115 to 0.165 ± 0.110 ($\Delta C = -0.022$), with a permutation-based $p < .0001$ that survived BH-FDR correction ($q < .05$). Furthermore, using weak FWER control, we identified statistically significant connectivity shifts for both VAS and VTS. During VAS, connectivity from the parietal regions (specifically the postcentral gyrus and the precuneus) to the frontal regions decreased in the beta band (13-30 Hz) compared to the pre-stimulation period (Fig. 4A-B). A similar pattern of decreased connectivity was observed after stimulation, though in a different anatomical location, specifically from the left calcarine to left cingulum (Fig. 4C). This effect was frequency-specific and consistently observed across multiple anterior-posterior IC pairs. Following VTS, we observed a significant increase in connectivity between the left superior medial frontal gyrus and the left orbital frontal region compared to the

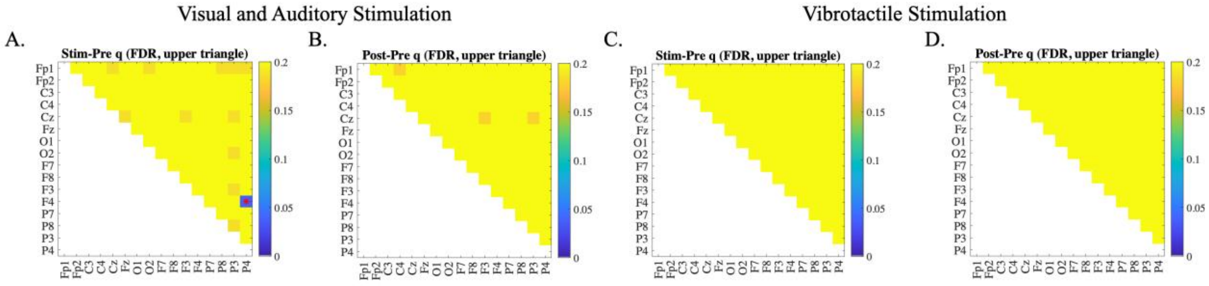


**Figure 3.** Channel-based coherence comparison between the stimulation and pre-stimulation periods in the gamma band, evaluated with Benjamini-Hochberg FDR correction (q values). The red dot highlights the only channel pair that reached statistical significance (F4-P4 in A). No statistically significant differences were observed in B, C and D. **A.** Coherence comparison between the stimulation and pre-stimulation period in the VAS condition. **B.** Coherence comparison between the post-stimulation and pre-stimulation period in the VAS condition. **C.** Coherence comparison between the stimulation and pre-stimulation period in the VTS condition. **D.** Coherence comparison between the post-stimulation and pre-stimulation period in the VTS condition.

pre-stimulation period (see Fig. 4D). This change in connectivity was observed in the gamma band, which corresponds to the 40 Hz stimulation frequency.

In Supplementary Fig. 1, we computed the event-related potentials for one subject during 2 minutes of VAS (26 samples per 100 ms epoch). We observed that among the seven EEG channels fluctuating at 40 Hz, two channels from the occipital region showed more than double the amplitude and were antiphase to the other channels. We found that 2 out of 5 participants had this approximate $\pi$ phase difference between the occipital and the other entrained cortices. We could not reliably estimate the phase difference in the remaining subjects because the sample rate was still too low.

We conducted verbal interviews with participants after undergoing the experiment. Their assessments of each stimulation modality are summarized in Table 1. Participants were asked to evaluate how long they could receive the stimulation daily for months and rate their tolerance on a scale from 0 to 10, where 0 indicated extreme difficulty in tolerating the stimulation and 10 indicated no difficulty. The tolerance scores for VAS and VTS were 6.2 ± 2.4 and 8.8 ± 1.3, respectively, showing higher tolerance for the VTS ($p$ = .00027). All participants indicated higher daily duration tolerance for VTS (67.7 min ± 75.8) compared to VAS (11.5 min ± 15.4; $p$ = .0049). Participants reported experiencing eyesight blurring from the visual stimulation, finding the auditory stimulation to be the most irritating sensory modality, and feeling bored during VAS. The consensus was that VTS is a more tolerable stimulation modality than VAS, causing less discomfort.

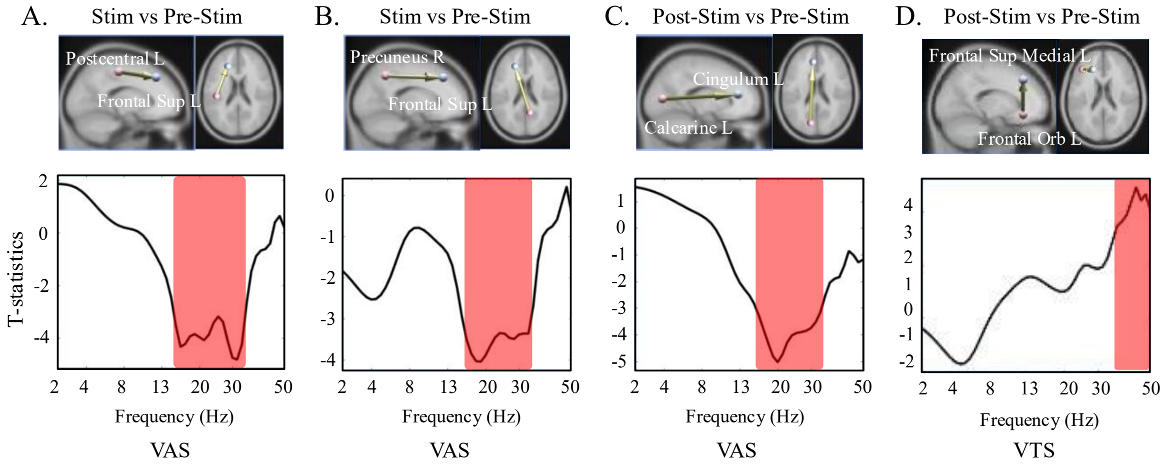


**Figure 4. A.** Connectivity shift measures with VAS (comparing the stimulation and pre-stimulation periods) in the 13-30 Hz frequency band between the left postcentral gyrus and left superior frontal cortex, depicted in a connectome connectivity map. The red-highlighted regions of the frequencies indicate t-statistics with an absolute value greater than 3 ($|t(14)| > 3$, $p < .01$). **B.** Same as A except the regions are the right precuneus and the left superior frontal cortex. **C.** Same as A except the comparisons are between the post-stimulation and pre-stimulation periods, and the regions are the left calcarine and left cingulum**. D.** Same as C except that it is for VTS, and the regions are the left superior medial frontal gyrus and the left orbital frontal region.

**Table 1. Participants verbal responses to the stimulation modality.**

| Stimulation Type | Visual and Auditory | Vibrotactile | P-value |
|---|---|---|---|
| Tolerance | 6.2 ± 2.4 | 8.8 ± 1.3 | < .01 |
| (0: intolerable / 10: tolerable) | | | |
| Daily duration tolerance (minutes) | 11.5 ± 15.4 | 67.7 ± 75.8 | < .01 |
| Preferred modality (15 subjects) | 1/15 | 14/15 | |

**Disclosure of interest.** No potential competing interest was reported by the author(s).

**Funding Information.**

Mazen Al Borno, Brice McConnell, Peter Teale, Coleman Institute for Cognitive Disorders

Mazen Al Borno, Office of Research Services, CU Denver

### Supplementary Material

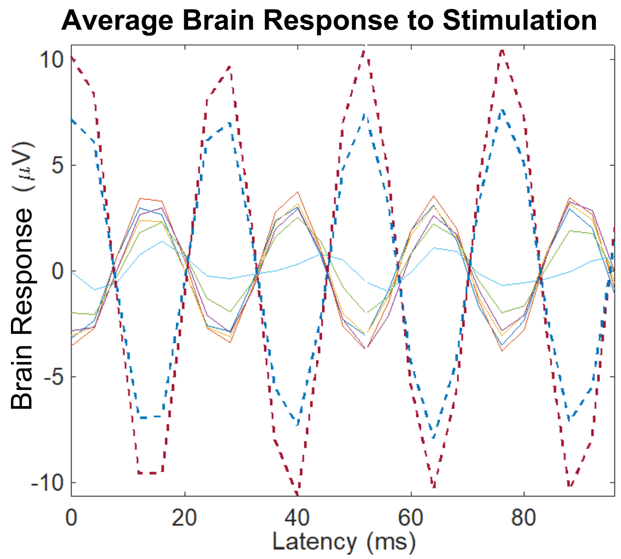


**Supplementary Figure 1.** Event-related potentials for one subject during VAS. Solid lines represent the prefrontal, frontal and central cortex and dotted lines represent the occipital cortex.
